# Improving Bacterial Genome Assembly Using a Test of Strand Orientation

**DOI:** 10.1101/2022.07.06.499059

**Authors:** Grant Greenberg, Ilan Shomorony

## Abstract

The complexity of genome assembly is due in large part to the presence of repeats. In particular, large reverse-complemented repeats can lead to incorrect inversions of large segments of the genome. To detect and correct such inversions in finished bacterial genomes, we propose a statistical test based on tetranucleotide frequency (TNF), which determines whether two segments from the same genome are of the same or opposite orientation. In most cases, the test neatly partitions the genome into two segments of roughly equal length with seemingly opposite orientations. This corresponds to the segments between the DNA replication origin and terminus, which were previously known to have distinct nucleotide compositions. We show that, in several cases where this balanced partition is not observed, the test identifies a potential inverted misassembly, which is validated by the presence of a reverse-complemented repeat at the boundaries of the inversion. After inverting the sequence between the repeat, the balance of the misassembled genome is restored. Our method identifies 31 potential misassemblies in the NCBI database, several of which are further supported by a reassembly of the read data.

## 1 Background

The study of bacterial function and interaction has the potential to revolutionize modern medicine through personalized treatment and pathogen discovery (Turnbaugh *et al.*, 2007; Le Chatelier *et al.*, 2013). In recent years, comparative genomics has greatly impacted our understanding of the diversity of bacteria. Moreover, due to advances in sequencing technologies, the number of bacterial genomes being sequenced grows at a fast pace, making it critical that we develop accurate and scalable analysis pipelines.

An important initial step in the analysis of genomic data is genome assembly, a complex and computationally intensive process that impacts any downstream application (Breitwieser *et al.*, 2017). The complexity of genome assembly arises in large part from repetitive sequences (repeats) in the genome. For short-read assemblers, exact repeats longer than readlength often cause highly segmented assemblies with many contiguous sequences (contigs) (Bankevich *et al.*, 2012; Zerbino and Birney, 2008). Furthermore, long, inexact repeats also pose a challenge for long-read assemblers since they can be difficult to distinguish given the higher sequencing error rates (Koren *et al.*, 2017; Vaser *et al.*, 2017; Kamath *et al.*, 2017; Kolmogorov *et al.*, 2019). Recent hybrid assembly methods, (Koren *et al.*, 2012; Haghshenas *et al.*, 2020) which combine short and long-read data, have overcome some of these challenges, but remain somewhat cost-prohibitive and limited in scope (Chin *et al.*, 2013).

Ambiguities in genome assembly are further compounded by reads coming from either strand of the original DNA sequence (Zeitouni *et al.*, 2010). If the genome contains long “inverted repeats” (i.e., repeats whose copies are on opposite strands), a misassembly can lead to inversions of very long segments of the genome. In this work, we investigate the effectiveness of *k*-mer frequencies in determining the relative orientation of genomic sequences. We demonstrate how this can be used for detecting and correcting these long incorrect inversions. Moreover, we explore how this idea can help prevent these misassemblies during the de novo assembly process.

It is well known that *k*-mer frequencies remain relatively constant throughout a prokaryotic genome for small *k* (Noble *et al.*, 1998; Mrázek, 2009; Pride *et al.*, 2003). In metagenomics, the study of the genomic content of microbial communities, the frequencies of *tetranucleotides* (sequences of four bases) are widely used (Kang *et al.*, 2015; Lu *et al.*, 2017); by grouping genomic sequences with similar tetranucleotide frequencies (TNF), metagenomic binning algorithms cluster assembled contigs that likely belong to the same genome. Figure 1 illustrates this phenomenon. The TNFs of several non-overlapping segments of different genomes are computed, and plotted using *t*SNE. We notice that segments from the same genome cluster together, and species from the same genus tend to have similar TNFs. Motivated by this remarkable usefulness of TNFs, we asked the following question: Could TNFs be used to distinguish the forward and reverse strands of a genome?

**Fig. 1.**
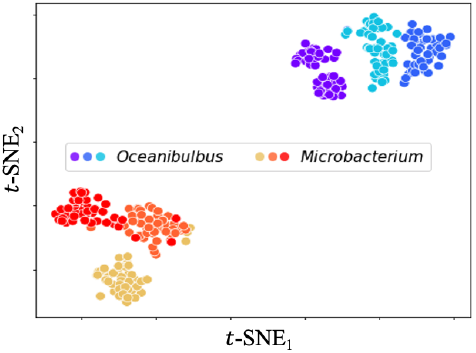
Two genera (Oceanibulbus and Microbacterium) and three species from each genus were chosen at random. Fifty non - overlapping segments, each with 50 kbp, were extracted from each of the six genomes. The oriented TNF was calculated for each segment, and *t*SNE was used to visualize the full set of TNFs. The plot shows both intra-genus and intra-species clustering.

In principle, one might not expect the TNF of both strands to be much different. In fact, metagenomics pipelines typically treat tetranucleotides and their reverse complement as the same tetranucleotide, giving rise to an “orientation-free” TNF vector with length 136, rather than 4^4^ = 256 (Kang *et al.*, 2015; Wu *et al.*, 2014). Nonetheless, our results surprisingly indicate that valuable information exists in the length-256 “oriented” TNF. In most cases, the TNF of the two strands of a genomic segment are markedly different, making it possible to determine the relative orientation of contigs. This observation naturally suggests that TNF can be used for repeat resolution on assembly graphs and for identifying inverted misassemblies on finished genomes.

In order to develop a mathematically sound TNF-based orientation test, we utilize a probabilistic model for generating a genome with a given TNF. This model is based on a framework previously used to study the information-theoretic limits of metagenomic binning (Greenberg and Shomorony, 2019). Note that an i.i.d. model is not capable of generating a sequence with a pre-specified TNF, since consecutive tetranucleotides in a sequence overlap by three bases. For this reason, we model the genome (in its forward orientation) as a third-order Markov model. Under this framework we derive a likelihood-based orientation test which uses the *entropy rate* as a way to decide whether two sequences are more likely to have the same orientation or opposite orientations.

We explore two main kinds of application of this test. First, in Section 2.3, we apply the test to many finished bacterial genomes from NCBI (Clark *et al.*, 2016). We find that in many cases, the genome can be cleanly partitioned into two sections that, from a TNF perspective, seem to have opposite orientations. Interestingly, this phenomenon arises as a result of DNA replication, which begins at a specific site in the genome called the *origin*, and proceeds bidirectionally until meeting at the *terminus* site. As previously reported, the direction of replication often has a nucleotide-composition (e.g., GC) skew (McLean *et al.*, 1998; Merrikh and Merrikh, 2018). This effectively divides the genome in two sections, typically in physical balance (i.e., equal size), with inverted TNFs (Song *et al.*, 2003).

In addition to the balanced two-section genomes, a small but significant proportion of the genomes examined had two parts of drastically different sizes. We postulate that in some of these “imbalanced” cases, a large erroneous inversion occurred during assembly. We look for long reverse-complemented repeats that may have caused the misassembly, and check if the physical balance of the genome is restored by inverting the sequence between the two copies of the repeat. Figure 2(a) depicts a heatmap of relative orientation along a real *S. enterica* genome from NCBI (National Center for Biotechnology Information, 1988). The heatmap clearly shows a significant imbalance in sizes of each region. In contrast, Figure 2(b) shows that, after inverting the sequence between a large repeat found in the genome, the heatmap is now balanced. We identify 31 examples of potential misassemblies on genomes from GenBank (Clark *et al.*, 2016) and NCTC 3000 (Public Health England *et al.*, 2014), each of which we correct using repeats in the genomes. In Section 2.4, we describe in more detail how the orientation test is used to correct inverted misassemblies.

**Fig. 2.**
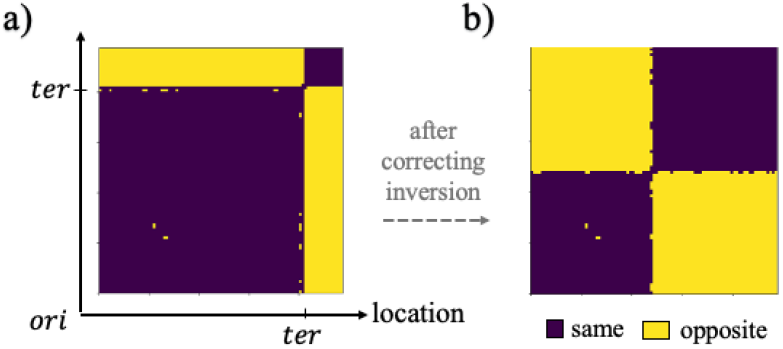
Correction of an Inverted Misassembly. (a) Heatmap of relative orientations along the genome is shown with labeled origin (ori) and terminus (ter) regions. The heatmap is clearly imbalanced as ori and ter are not evenly spaced. (b) Heatmap of corrected genome is balanced after inverting the sequence between a long repeat found.

In Section 2.5, we explore the second use of the TNF orientation test by resolving ambiguities in genome assembly. In particular, we consider the resolution of assembly graph structures caused by repeats by comparing the orientation between incoming and outgoing nodes of the repeat node. We focus on paired-end read datasets, and use the paired-end information only to provide us with a ground truth for the node resolution step. We find that, in nearly 75% of cases where the TNF orientation information can be used to resolve a node, it suggests the correct resolution. While this accuracy is below what would be desirable to reliably resolve repeat nodes in assembly graphs, it shows the test provides useful information, which could be used in conjunction with, say, paired-end or Hi-C data (Lieberman-Aiden *et al.*, 2009).

Overall, this work explores a promising new use of TNF (and, in general, *k*-mer composition): disambiguating strand orientation. In addition to being useful for improving assembly, this idea could also find applications in the study of genomic rearrangements in bacteria. The code to analyze strand orientation on assembled genomes, locate origin and terminus regions, and correct potential misassemblies is available at https://github.com/gcgreenberg/Oriented-TNF.git.

## 2 Results

In order to explore the use of TNF to detect strand orientation, we propose a likelihood-based test. In the next subsection we present the test, but defer the description of the model and test derivation to Section 3. In Sections 2.2-2.4, we use the test to create a heatmap of relative orientations which is then used to find and correct an inverted misassembly. In Section 2.5 we explore results of the test in resolving genome assembly graph structures.

### 2.1 TNF-based Orientation Test

Given two sequences from the same genome, **x** and **y**, the test *δ* decides whether **y** has the same or opposite orientation as **x**. The test is as follows:

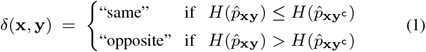

Here, 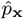 represents the empirical (third-order) Markov model obtained from a sequence **x**, and *H* is the *entropy rate* of a Markov process (Cover and Thomas, 2006), both derived in Section 3.1. We also let **xy** denote the concatenation of **x** and **y**, and **x**^c^ denote the reverse-complement of **x** (e.g., (ATTC)^c^ =GAAT). It is important to note that *δ* only relies on the TNFs of **x** and **y** since the entropy rate can be written in terms of the TNF only (see Section 3.1). Intuitively, a lower entropy rate signifies a less “random” sequence. In general, we expect that, if **x** and **y** have the same orientation, **x** and **y**^c^ will be mismatched, increasing the value of 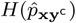 and making the test more likely to output “same.”

### 2.2 Orientation Matrix

We first use the test in Eq. (1) to visualize the relative orientations of windowed segments of a genome. The length of each window was chosen to be 100kbp, with a stride (i.e., distance between the start of consecutive windows) of 50kbp. The window length was chosen to be as small as possible while ensuring the orientation test is robust to local fluctuation in TNF. For each extracted window, we calculate the TNF of the genome segment. The orientation test is then performed for each pair of windows using the calculated TNFs. The result is a matrix of orientation tests as shown in the heatmap of Figure 3(c).

**Fig. 3.**
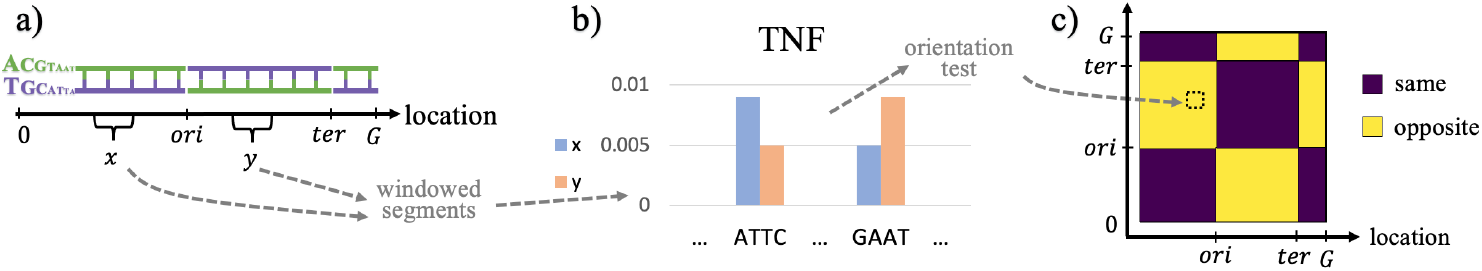
Computation of Orientation Matrix. (a) A bacteria genome with labeled replication regions, ori and ter. (b) Orientation test between windowed segments of the genome, **x** and **y**, represented as a comparison of their TNFs. In this case, reverse-complement tetranucleotides, ATTC and GAAT, have opposite frequencies in **x** and **y**, suggesting that *δ*(**x**, **y**) =“opposite”. (c) The orientation matrix with two clear sections of opposite orientation.

Next, we use the orientation matrix to locate the DNA replication origin (*ori*) and terminus (*ter*). As discussed in Section 1, the replication regions generally divide abacterial genome in two sections of opposite orientation. With respect to Figure 3(c), the windows which contain *ori* and *ter* likely lie at the apparent transitions in orientation. To locate the transitions, a clustering method is used to group windows on the orientation matrix. Our results indicate that a simple spectral method (using the first principal component of the orientation matrix) effectively pinpoints the transitions. In Section 3.3, we provide an algorithm to obtain precise estimates for the locations of the two replication regions.

### 2.3 Analysis of Assembled Genomes

We randomly chose 5,000 genomes out of the collection of over 22,000 completed genomes in NCBI’s GenBank (Clark *et al.*, 2016). On each chosen genome, we aim to: discover any irregularities in the oriented TNF along the genome; identify the origin and terminus sites of replication; and algorithmically locate and correct inverted misassemblies (Section 2.4).

The binary heatmaps of Figures 4(a-e) depict the orientation matrices for five chosen genomes with common patterns. Figures 4(a-d) depict examples of “balanced” genomes; i.e., genomes divided in two sections of roughly equal length. The transitions in orientation between these sections correspond to the origin and terminus regions of replication. In some cases (including Figures 4(a,b)), the genome is purposefully assembled so that *ori* lies at the start of the sequence (Kono *et al.*, 2017), which is justified since bacterial genomes are circular. It is also important to note that a simple rotation of the genomes in Figures 4(c,d) would accomplish the same task of producing heatmaps with only one apparent transition.

**Fig. 4.**
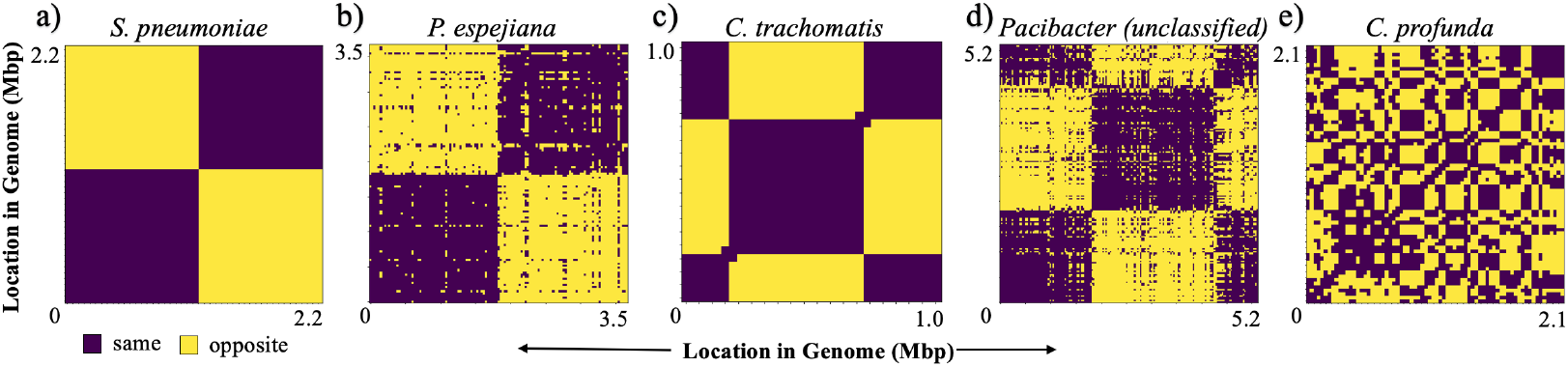
Common Orientation Matrix Heatmaps. (a-d) Genomes with two roughly equal sections of opposite orientation corresponding to the ori-ter axis. The heatmaps in (b) and (d) are noisier, indicating a weaker nucleotide-composition skew. (e) A genome lacking any clear partition and no strand-specific information. Each category of heatmap pattern shown is considered typical, in contrast to those in Section 2.4.

Notice that the orientation matrices in Figures 4(b,d) are much noisier than those of Figures 4(a,c), with many “opposite” orientations scattered throughout the matrix. In contrast, the heatmap of Figure 4(e) also contains considerable noise, but does not contain any clear transitions in orientation corresponding to the replication sites. This type of pattern was seen in a significant, but surprisingly small fraction of the genomes analyzed.

### 2.4 Correction of Inverted Misassemblies

In addition to the balanced orientation matrices shown in Figures 4(a-d), a number of genomes had two sections of highly uneven lengths, which we refer to as “imbalanced” genomes. Such a genome is unnatural, as the *ori* and *ter* regions typically lie opposite each other on the circular chromosome to optimize the efficiency of DNA replication (Song *et al.*, 2003). For this reason, we hypothesize that (some, if not all) imbalanced genomes contain an assembly mistake in the form of an inversion. In general, such a misassembly is due to a reverse-complemented repeat (i.e., a repeat on opposite strands): when attempting to extract a contig by traversing the assembly graph, the incorrect direction may have been chosen after the repeat node, creating an erroneous inversion.

In order to classify a genome, we define a measure of balance whose value increases the closer the two sections are in length (i.e., closer to 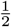 *G*).

#### Definition 1.

*Given a genome* 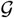 *of length G, and ori* < *ter locations, the **balance** is defined as*

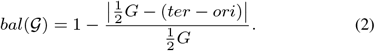

For each chosen GenBank genome 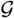, we first locate the replication sites. We consider 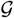 for misassembly detection if the method returns a clearly defined *ori* and *ter*, and if 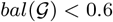. Next, using the NUCmer pipeline from MUMmer version 4.0 (Kurtz *et al.*, 2004), we search for a list of candidate repeats which may have caused the misassembly. If there exists a repeat 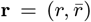 which satisfies all the following criteria, we claim that **r** is the source of the misassembly:

i. *r* and 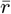 are reverse-complemented.
ii. The length of the repeat is at least 10kbp.
iii. The repeat has a minimum of 95% nucleotide identity.
iv. The locations of *r* and 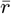 are on opposite sections of orientation in 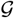.
v. If we create a new genome 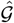 by inverting (i.e., reverse complementing) the sequence between *r* and 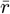, the “corrected” genome 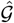 must have 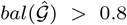.

In all, our method detected 31 potential misassemblies (with imbalance and inverted repeats *r* and 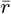) out of the 5,000 genomes analyzed. Of the 31, only six had complete read data available (see Supplementary Table T1). In Figures 5 and 6, we discuss the results from two such genomes.

**Fig. 5.**
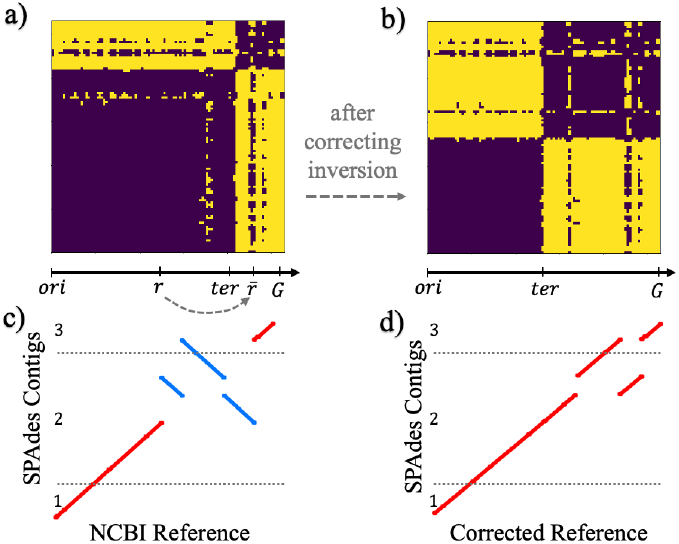
E. coli Strain A1_136 with Potential Misassemblies. (a) Orientation heatmap of original GenBank assembly is imbalanced (*bal* = 0.46). A long reverse-complemented repeat, *r*, 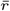 is used to correct the misassembly. (b) Orientation heatmap of the corrected genome is balanced, with evenly spaced replication regions. (c) Dot plot of the original genome compared with a new assembly of the read data using the SPAdes assembler. The SPAdes contigs are identical to the reference except for a large inverted sequence corresponding to the segment between *r* and 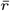. (d) Dot plot after correcting the misassembly in the original genome. The SPAdes contigs are nearly identical to the corrected genome (except for a small translated sequence), indicating the misassembly is properly corrected.

**Fig. 6.**
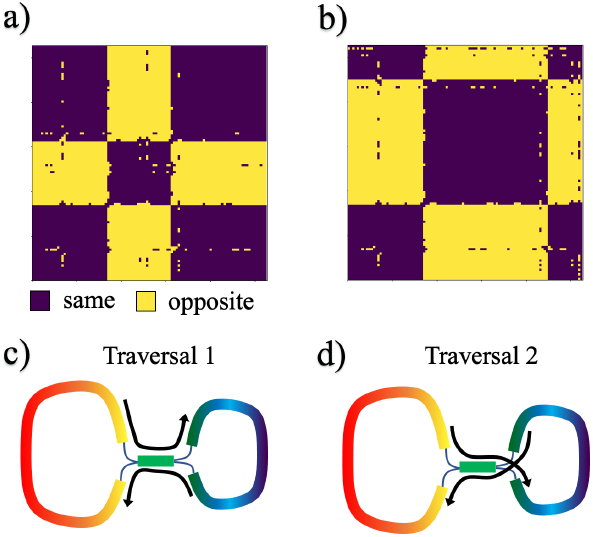
E. coli Strain RM9088 with Potential Misassemblies. (a) Orientation heatmap of GenBank genome is imbalanced (*bal* = 0.54). (b) Heatmap of corrected genome is now balanced. (c,d) Assembly graph using the HINGE assembler offers two possible traversals for completed genomes. Traversal in (c) corresponds to original genome and (d) corresponds to the corrected genome, which indicates that Traversal 2 is the correct one.

Figure 5(a) shows the orientation heatmap of an *E. coli* strain from GenBank (read data accession numbers SRR8549120 and SRR8549113), a clear example of an imbalanced genome. Using the process described above, we located a long repeat (11kbp) on either side of ter, as shown in the axis below the heatmap. After correcting the supposed misassembly by re-inverting the sequence between *r* and 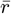, the genome becomes balanced (Figure 5(b)). Next, to provide additional evidence that this is indeed a misassebmly, we re-assemble the hybrid read data (i.e., containing both short and long reads) using the SPAdes assembler version 3.15.0 (Bankevich *et al.*, 2012). In Figure 5(c), we see that running SPAdes leads to an assembly identical to the NCBI reference, but with a large inversion with ends precisely where our method discovered the large repeat. After correcting the inversion, the resulting genome in Figure 5(d) is in agreement with the SPAdes assembly (except for a small non-inverted translation which is undetectable using the heatmap alone). Together, the balance of the corrected genome and the agreement of the SPAdes assembly provide strong evidence of a misassembly.

Next, we analyze a different *E. coli* strain (accession SRR9953605) that also has an imbalanced heatmap, shown in Figure 6(a). A long repeat of length nearly 20kbp was found satisfying the required criteria, and after correction we obtain the balanced genome in Figure 6(b). As in the genome of Figure 5, we wish to support our conclusion by re-assembling the genome. We utilize the HINGE assembler to assemble the long-read PacBio data, since HINGE attempts to produce a graph that captures all possible assemblies that are consistent with the data (Kamath *et al.*, 2017). Applied to this PacBio dataset, this results in a circular assembly with two possible traversals, as shown in Figures 6(c,d). As it turns out, Traversals 1 and 2 correspond nearly identically to the original and corrected version of the NCBI genome, respectively. Thus, the orientation heatmap indicates that Traversal 2 is likely the correct one. This way, our method allows one to resolve a long repeat in a principled manner during assembly.

In addition to the genomes of Figures 5 and 6, we assemble the read data from the other six GenBank misassemblies found. We also included an example from the NCTC database (Public Health England *et al.*, 2014), whose genome was known (from the supplemental information in Kamath *et al.* (2017)) to have an assembly similar to Figures 6(c,d). In most cases, including the genome of Figure 6, this led to an unfinished assembly due to a long inverted repeat (see Supplementary Figures S1-S3 for more details). The use of TNF orientation provides a novel method to complete such assemblies. In Section 2.5, we explore the use of sequence orientation to resolve more general assembly graph structures.

### 2.5 Resolving Assembly Graphs

The first step of most graph-based assemblers (particularly, for short reads) is to construct the de Bruijn graph by breaking reads into *k*-mers, for a specified *k*. Initially, each unique *k*-mer represents an edge between (*k* – 1)-mer nodes on the graph (Pevzner *et al.*, 2001). The de Bruijn graph is then simplified by merging unambiguous paths, and by resolving repeat nodes that are bridged by reads. Paired-end read information can be used to resolve additional nodes, and the resulting paths are extracted as contigs (Bankevich *et al.*, 2012).

Consider the graph structure shown in Figure 7(a), in which a repeat node **r** has two incoming edges, **w** and **x**, and two outgoing edges, **y** and **z**. Several ambiguities exist in such a structure. For instance, when attempting to create a contig traversing **w**, both **y** and **z** are potential options after **r**. Figure 7 illustrates how the orientation test can be applied to help disambiguate such graph structures. The TNF of each node is computed and, for each pair of incoming to outgoing nodes, {**w**, **x**} × {**y**, **z**}, the orientation test is performed. This produces a table of tests which determines the “compatibility” of each pair. Consider the table of Figure 7(c). The incompatibility (i.e., “opposite” orientation) of **x** and **y** implies not only that **y** does not directly follow **x** in the genome, but also that the correct contigs are (… **wry** …) and (… **xrz** …). The three other orientation tests are performed to verify this implication; if all are consistent with the first test (as is the case in Figure 7(c)), we label the table of tests as conclusive. If one or more of the four tests is inconsistent with the others (e.g., **w** and **y** are compatible, but **x** and **z** are not), then the table is inconclusive, and cannot reliably be used to determine the ordering of nodes. Similarly, if all pairs of nodes are compatible, then the table is indeed consistent but still inconclusive, since either pair of incoming and outgoing nodes are still possible.

**Fig. 7.**
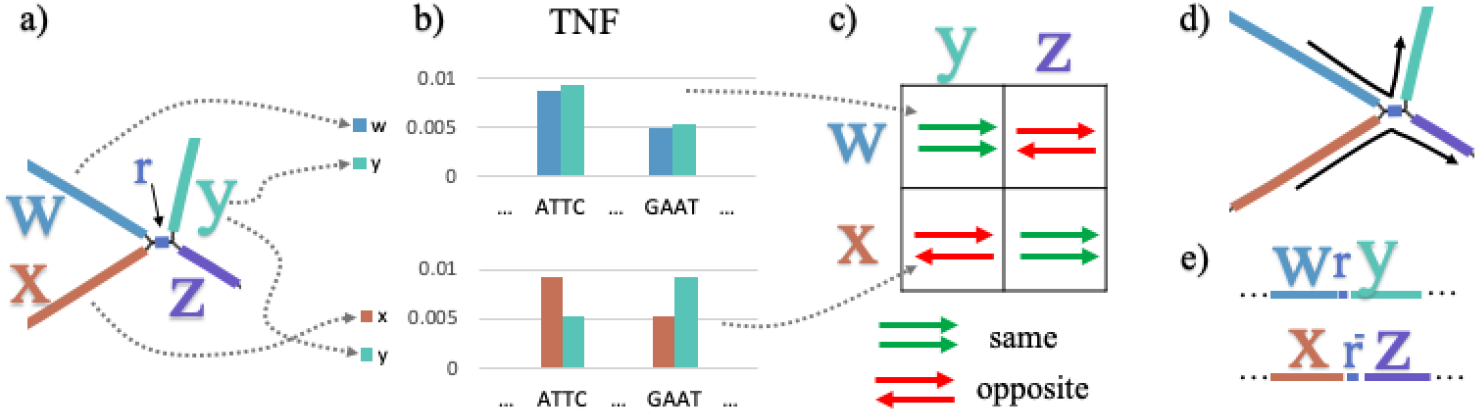
Pipeline to Resolve Repeats. (a) Assembly graph structure around repeat node **r** with incoming nodes **w** and **x** and outgoing nodes **y** and **z**. (b) Orientation tests between **w** and **y** (top), and between **x** and **y** (bottom) represented as a comparison of the TNFs of each node. (c) Orientation table of all pairwise tests between incoming and outgoing nodes. In this case, the table is conclusive since the the outcomes are consistent with a unique traversal of the graph shown in (d) and the corresponding contigs shown in (e).

We expect a conclusive table only under the following circumstances: (1) the repeat is reverse-complemented *and* both copies lie in the same orientation section or (2) the repeat is on the same strand *and* lies on the opposite sides of *ori* and ter. While this limits the scope of applicability of the method, it is important to note that it is complementary to the method in Section 2.4, which requires a reverse-complemented repeat on opposite sides of *ori* and *ter*.

Given a conclusive test on a repeat node which is not already traversed by any contigs in the assembly, contigs on either side of the repeat can be connected in the manner the test indicates. This procedure can also be generalized to repeat nodes with multiplicities greater than two.

We downloaded 15 paired-end read datasets from NCBI’s Sequence Read Archive (SRA) (Leinonen *et al.*, 2011) to evaluate the proposed method. Each is obtained from a distinct bacterial species, chosen randomly from a list of isolated and sequenced bacteria on SRA. We use SPAdes to assemble the datasets, resulting in two main outputs: an assembly graph (consisting of node sequences and overlap edges), and a set of contigs corresponding to paths on the graph. Repeat nodes that can be resolved by paired-end reads are left unresolved in the assembly graph. This way, we can perform the TNF-based node resolution on the graph and use the SPAdes contigs (obtained using paired-end read information) as ground truth to evaluate the accuracy of the orientation test.

We evaluate accuracy based on the two metrics shown in Table 1. The first metric measures the accuracy of the orientation test. Considering the graph of Figure 7(a), if (… **wry**…) is a ground-truth contig, for example, we perform the orientation test on **w** and **y**. If *δ*(**w**, **y**) = “same”, the test is deemed correct. The evaluation is performed on all ground-truth contigs in the dataset.

The second metric is the accuracy of the table. We a priori consider only the cases in which the table of orientation tests for a given repeat node is conclusive, allowing the repeat to be resolved. For a given repeat node **r**, if all contigs which traverse **r** match the table conclusion, then the table is considered correct. Conversely, if any contigs contradict the table, it is incorrect. For example, suppose the table concludes that the correct contigs should be (… **wry** …) and (… **xrz** …). If in reality, there are two ground-truth contigs traversing **r**, (… **wrz** …) and (… **xry** …), the test is incorrect. On the other hand, if there is only one associated contig, (… **xrz** …), then the table is deemed correct.

From Table 1, we see that there were only a small number of conclusive tests for each assembly. There were also a similar number of repeat nodes which would allow the method to resolve the repeat, i.e., with both a conclusive orientation table and no ground-truth contigs. It should be noted that for these nodes the method frequently connected long contigs to each other, creating a significantly more contiguous assembly. Nonetheless, considerable practical improvements are necessary before the method can be reliably used to resolve repeats. We discuss some directions for further improvement in Section 4.

**Table 1.**
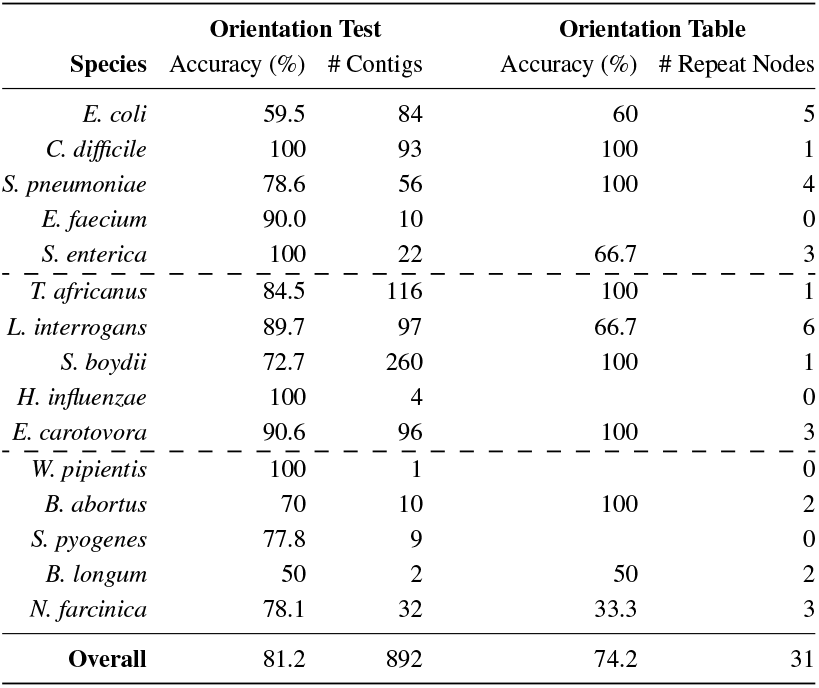
Accuracy assessment for Orientation Test. Contigs containing repeat nodes provide ground truths for the method. The Orientation Test column measures the accuracy of the test on all individual contigs which span a repeat node. The orientation test is performed on the sequences before and after the repeat and if the result is “same”, the test is considered correct. The Orientation Table column measures the accuracy of the table of orientation tests in resolving repeat node graph structures. For each repeat node **r** with one or more contigs spanning it, the table of tests is computed. A table is considered correct if all contigs containing **r** are consistent with the conclusion of the table. If the table is inconclusive, it is discarded.

## 3 Methods

In order to derive the orientation test of Eq. (1), we model a bacterial genome as a third-order Markov process *p*. In this section, we present this model in detail and discuss its effectiveness.

### 3.1 Markov Model and Properties

Given the alphabet *χ* = {*A,T,G,C*}, let **a** ∈ *χ*^3^ and *b* ∈ *χ*. A third-order Markov process is defined by an initial state distribution and a transition probability matrix *p*(*b*|**a**). The third- and fourth-order steadystate distributions, *p*(**a**) and *p*(**a***b*), can be uniquely derived from the transition probabilities. The TNF of a genome can be thought of as its fourth-order steady state distribution (under the natural assumption that the Markov process begins in steady state). Note that the symbol *p* is used for each distribution, and can be distinguished by the argument.

We first define two information-theoretic measures (Moulin and Veeravalli, 2018; Cover and Thomas, 2006; Greenberg and Shomorony, 2019) used in the properties following.

#### Definition 2.

*The KL divergence rate between p and q is given by*

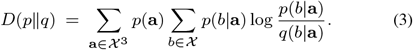

#### Definition 3.

*The entropy rate of p is given by*

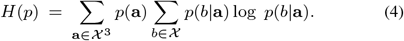

Suppose *p* generates a genomic sequence **x** = *x*_0_*x*_1_ … *x_L_–1*. The *empirical* TNF of **x**, 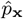, can be determined simply by calculating the frequency of each four-letter combination in the sequence. A straightforward calculation yields the following result.

#### Proposition.

*The probability of* **x** *under p is given by*

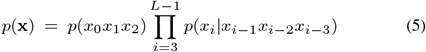

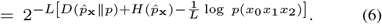

Notice that, as the length *L* increases, the effect of the initial state becomes negligible, as suggested by the 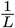 coefficient in Eq. (6). Formally, the limit of the *normalized log-likelihood* of **x** can be written as

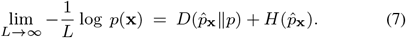

Suppose now that the distribution which generated **x** is unknown. We may wish to determine the distribution from **x** itself. In fact, the maximum-likelihood (ML) distribution — the distribution under which **x** has the highest likelihood — is 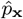 itself (Cover and Thomas, 2006). Concretely, the ML probability of **x** is given by

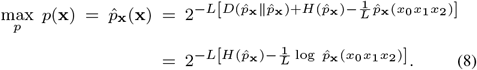

Notice that Eq. (8) is identical to Eq. (6) without the *D* term. Consequently, the normalized log likelihood under the ML distribution 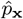 is simply the entropy rate, 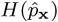.

### 3.2 Orientation Test

We use the Markov model described above to create a test of strand orientation. We consider the scenario in which we have two sequences from the same genome, **x** and **y**, and construct a hypothesis test for whether **x** and **y** are on the same strand. We decide between the two hypotheses

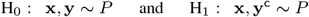

where *P* is the generating distribution. The prior probabilities of H_0_ and H_1_ are *π*_0_ and *π*_1_. The prior distribution on *P*, however, is unknown and in general, the genome of origin cannot be used to determine *P*. Thus, we use the ML probabilities to choose the correct hypothesis, resulting in a *generalized likelihood ratio test* (GLRT) (Moulin and Veeravalli, 2018). Assuming that *L* is large enough that the effect of the initial state is negligible in Eq. (8), the resulting test is

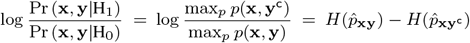

where **xy** and **xy**^c^ represent the corresponding sequences concatenated. Moreover, the decision rule *δ* for the binary hypothesis test that minimizes the error probability is the following:

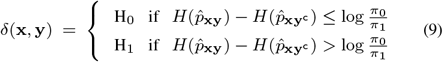

The GLRT in Eq. (9) tests the hypothesis that two sequences from the same genome have the same relative orientation. In Sections 2.3 and 2.4, we used the test to detect replication sites and inverted misassemblies, and in Section 2.5 we disambiguate genome assembly graph structures. In either case, we set the priors equal 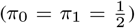. This is justified for completed genomes since the division caused by replication sites makes “same” and “opposite” equally likely, and in assembly, contigs are equally likely to lie on either strand of the chromosome.

### 3.3 Estimating Replication Regions

In Section 2.2, we use the method introduced above to determine the orientation matrix of a genome. Algorithm 1 uses the matrix to obtain a precise estimate of *ori* and *ter* sought in Section 2.3, as a function of window length and stride. Specifically, we choose the center of the range of potential locations, which we motivate in the following remark.

Let 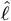 be the start of the window on the left side of a transition in orientation, ℓ be the true location of a transition (i.e., ori or ter), *W* be the window length, and S be the stride.

#### Remark.

The potential range of locations for the transition is

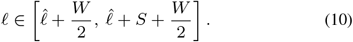

#### Explanation.

Suppose **w**_0_ is a window to the left of and away from the transition, and **w** is a second window which may overlap with the section of opposite orientation in some proportion. Let us make the following assumption: in order for the orientation test to decide **w** is of opposite orientation relative to **w**_0_, at least half of **w** must lie in the opposite section. The above assumption allows us to conclude that at most half of the window to the left of the transition can be past ℓ. In other words 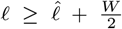. Moreover, the converse assumption that *δ*(**w**, **w**_0_) = “same” when at most half of **w** is past the transition point indicates that at least half of the right window must be past ℓ. This leads to 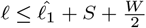.

**Algorithm 1:**
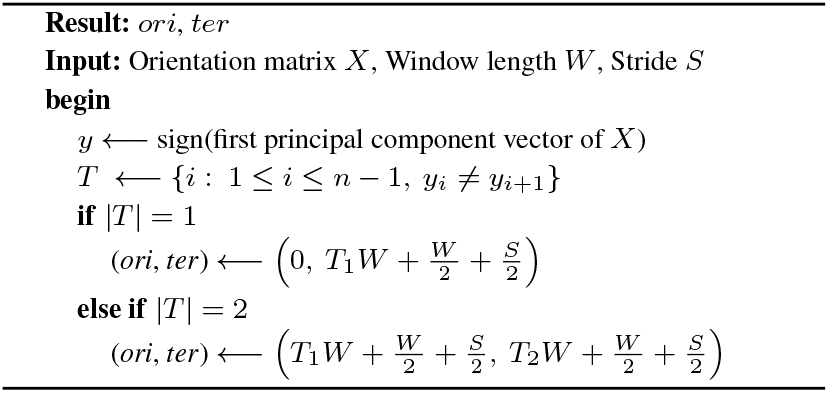
Estimating the locations of *ori* and *ter*

## 4 Discussion and Future Work

In this work, we explored the use of TNF to improve bacterial genome assemblies. Based on a model of genomic sequences, we derived a test of sequence orientation and with it, proposed two distinct applications: a method to correct misassembled genomes and a procedure to resolve repeats during assembly.

We point out that, in our approach to looking for inversion misassemblies, we sought to identify inversions between repeats that lie in sections of opposite orientation. However, in principle we do not need to restrict our search those types of inversion. Suppose the genome has a repeat with both copies in the same orientation section. An erroneous inversion between the two copies of this repeat would create a visible inversion in the heatmap (secondary to the “natural inversion” present from the replication skew), making the genome appear to have more than two orientation sections. Hence, one could hope to identify such misassemblies by simply counting the number of orientation transitions. While we did discover several examples where a secondary inversion is interposed in an otherwise balanced genome, in all the cases we identified, the read data did not suggest that what we had was in fact a misassembly. This may be because the identified inversions were in fact real biological artifacts (e.g., horizontal gene transfer or chromosomal rearrangement).

Another possible misassembly scenario involves an incorrectly translated segment, which can be caused by a triple repeat of the same orientation in the genome (see Supplementary Figure S4). Depending on the locations of the repeat, such a translation would be visible in the heatmap. A future research direction could be to expand our method to correct translated misassemblies, or even to resolve the triple repeat in the first place during the assembly process.

In the context of repeat node resolution in assembly graphs, the accuracy results described in Section 2.5 are below what would be needed for the orientation test to be incorporated into an assembly pipeline. To improve on our proposed methods, we require a deeper understanding of the effects of the “natural inversion” due to the replication sites. The low accuracy could also be due to the fact that, in some genomes, the TNF of the two DNA strands are not markedly distinct, and the test should not be used. One direction for future work is to determine which global TNF distributions make the TNF-based orientation test not reliable.

Beyond the orientation test itself, we believe promising applications of the oriented TNF exist for metagenomics methods. For instance, in metagenomic binning, the TNF in general is rendered orientation-free by combining reverse-complemented tetranucleotides. However, as evidenced in our results, the oriented TNF may provide more specificity in distinguishing contigs. Another viable application is metagenomic assembly, the highly complex process of assembling many species’ genomes concurrently using metagenomic data. In metagenomic communities, repeats are often shared across species (Nurk *et al.*, 2017), meaning that for the graph in Figure 7(a), **w** could be a sequence from an entirely different species than **z**, for example. In these cases, we could expand the repeat resolution method to not only determine strand orientation, but also differentiate between species.

