## Supplementary Information for "Improving Bacterial Genome Assembly Using a Test of Strand Orientation"

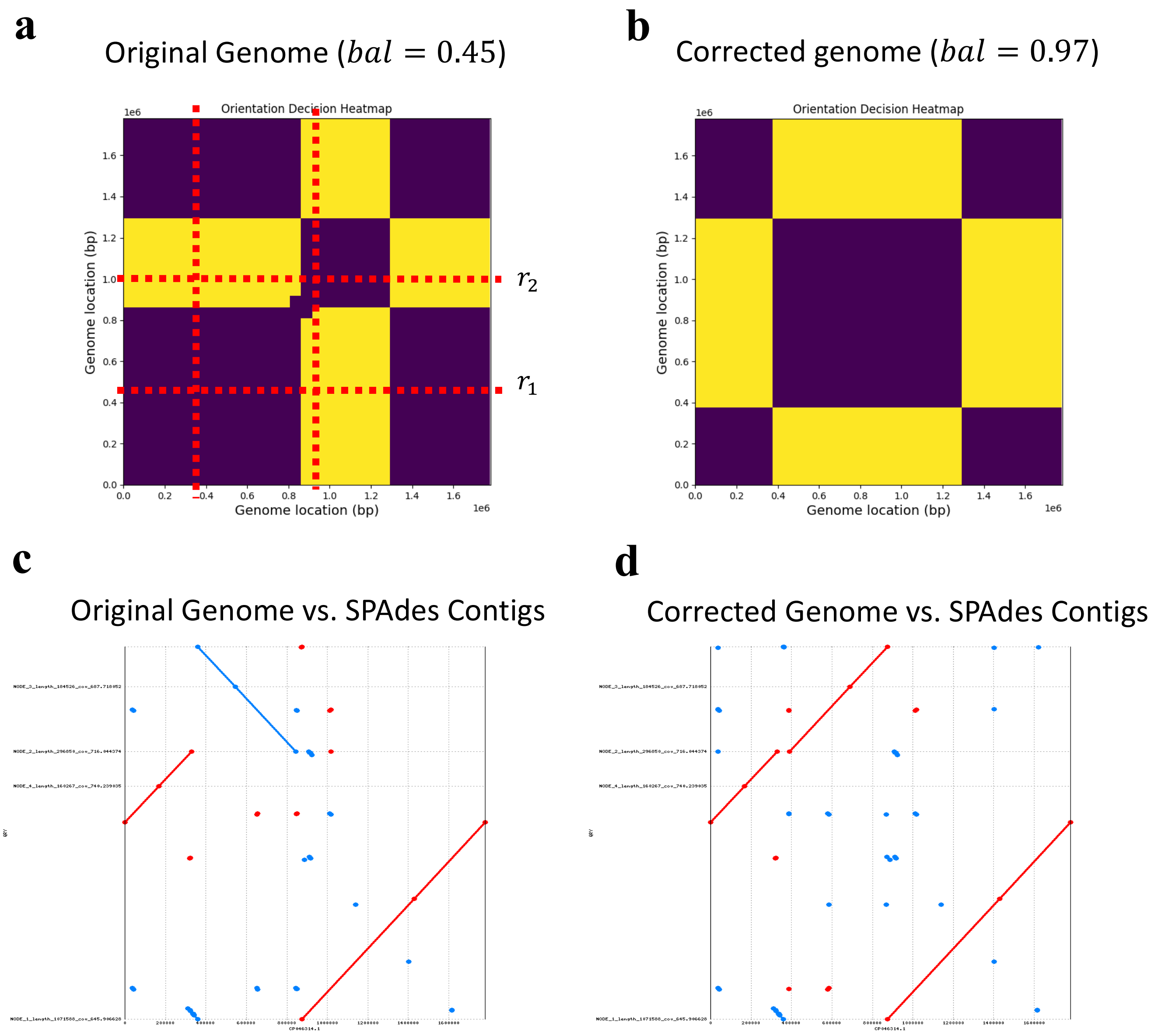

**Supplementary Figure S1.** Correction of Inverted Misassembly of *G. morbillorum* (strain: FDAARGOS_741; taxID: 29391; assembly accession: GCA_009730315.1; BioSample: SAMN11056456). **(a)** Orientation heatmap of original genome from GenBank is highly unbalanced. Red dashed lines represent the locations of the repeat $(r_1, r_2)$ used to correct the misassembly. **(b)** Heatmap of corrected genome is now balanced. **(c)** Dot plot of original genome against largest four contigs from the re-assembly of the hybrid read data using SPAdes assembler. A large inversion is manually positioned to illustrate the location of the repeat. **(d)** Dot plot of corrected genome against SPAdes re-assembly. The large inversion from (c) is now un-inverted, indicating that the ordering of the contigs chosen in (c,d) is indeed correct.

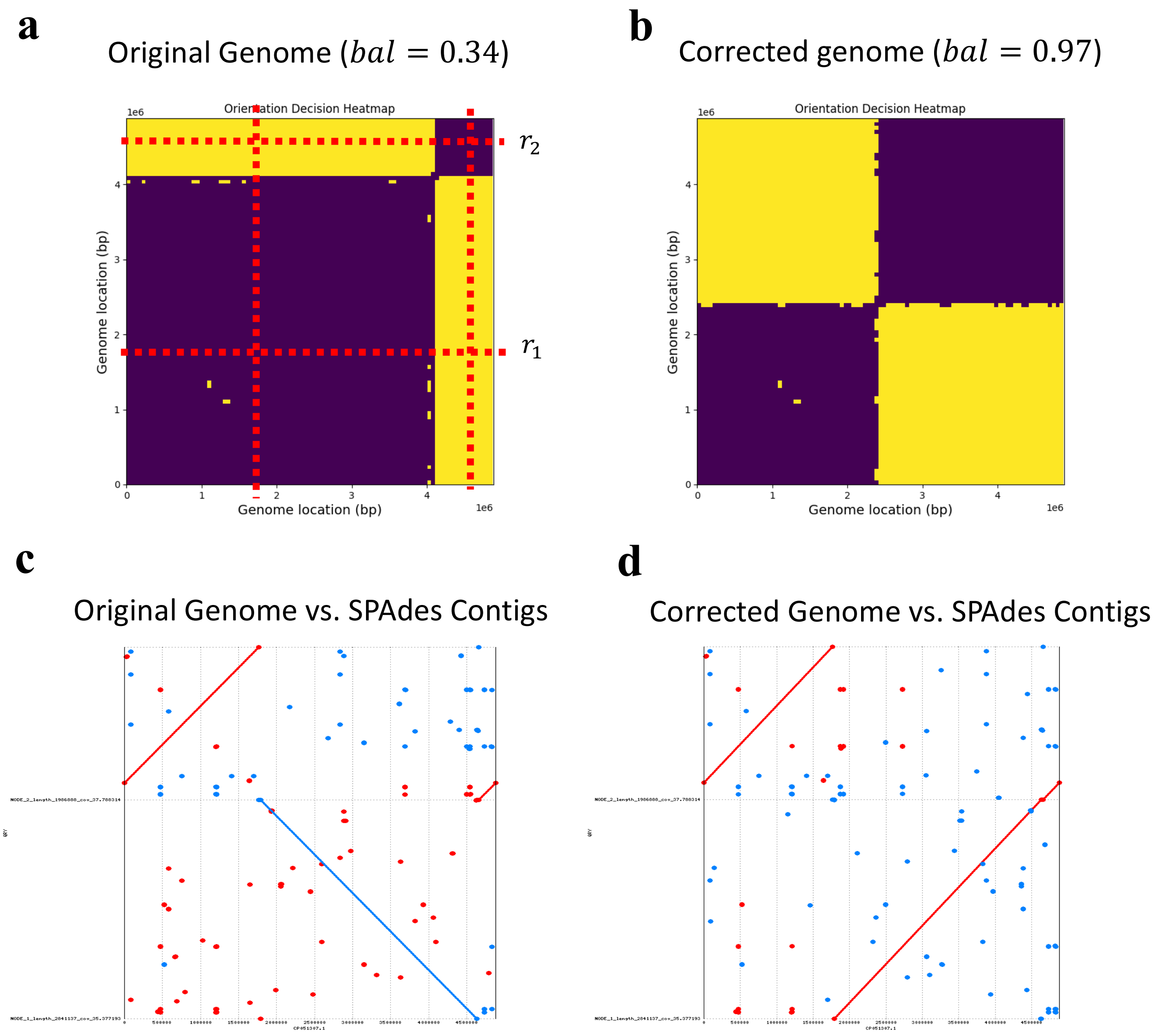

**Supplementary Figure S2.** Correction of Inverted Misassembly of *S. enterica* (strain: CVM 35189; taxID: 28901; assembly accession: **GCA_016451985**.1; BioSample: SAMN14504941). **(a)** Orientation heatmap of original genome from GenBank is highly unbalanced. Red dashed lines represent the locations of the repeat ($r_1, r_2$) used to correct the misassembly. **(b)** Heatmap of corrected genome is now balanced. **(c)** Dot plot of original genome against largest four contigs from the re-assembly of the hybrid read data using SPAdes assembler. A large inversion is manually positioned to illustrate the location of the repeat. **(d)** Dot plot of corrected genome against SPAdes re-assembly. The large inversion from (c) is now un-inverted, indicating that the ordering of the contigs chosen in (c,d) is indeed correct.

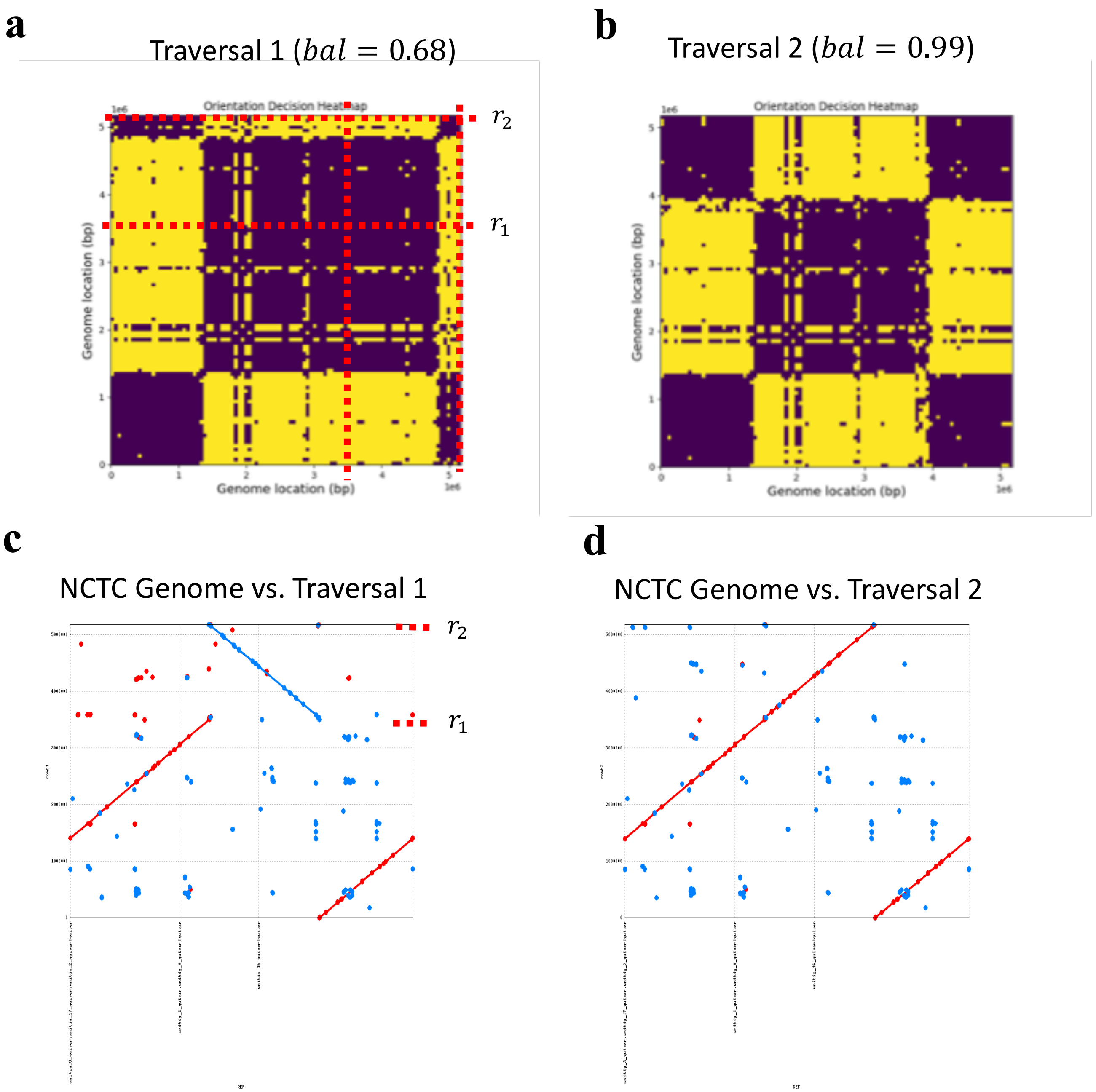

**Supplementary Figure S3.** Assembly analysis of *E. coli* (strain: NCTC9006; taxID: 562; BioSample: SAMEA3376915). The long read data from NCTC 3000 was assembled using the HINGE assembler, resulting in an assembly graph with two possible traversals. **(a)** Orientation heatmap of Traversal 1 is unbalanced. Red dashed lines represent the locations of the repeat ($r_1, r_2$) which causes the incomplete assembly. **(b)** Heatmap of Traversal 2 is balanced, suggesting that it is the correct assembly of the genome **(c)** Dot plot of the NCTC genome (containing three contigs) against Traversal 1. A large inversion is present exactly between ($r_1, r_2$). **(d)** Dot plot of NCTC genome against Traversal 2. The large inversion from (c) is now un-inverted, indicating that the ordering of the contigs chosen in (c,d) is indeed correct.

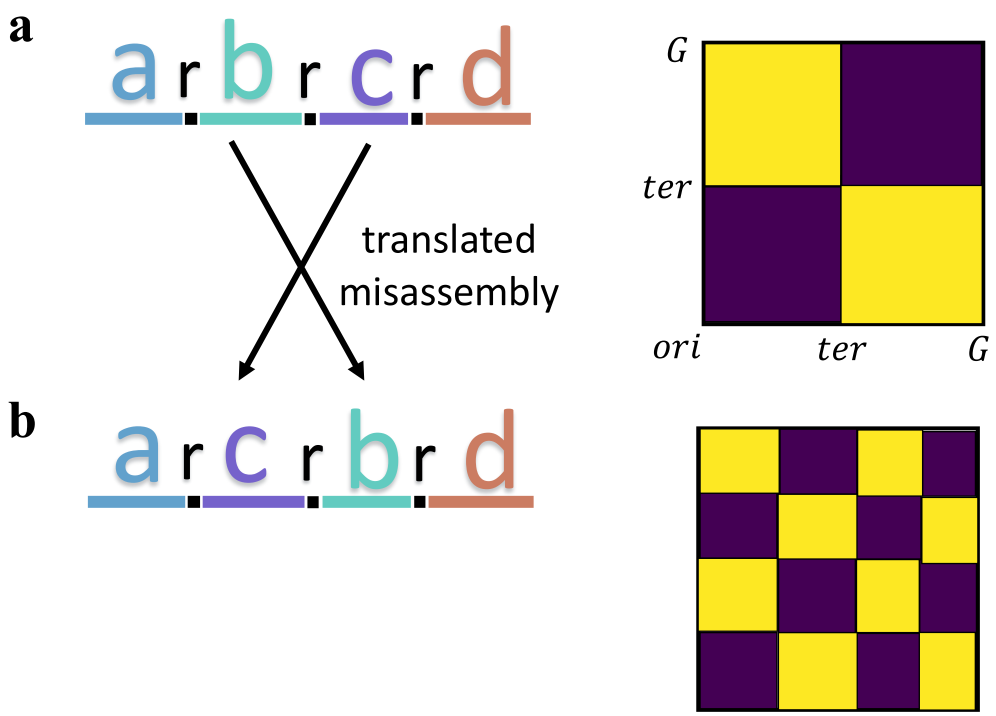

**Supplementary Figure S4.** **(a)** Example of a genome with a triple repeat on the forward strand, along with the corresponding orientation heatmap. **(b)** Depiction of the same genome if an erroneous translation occurred during assembly by switching the locations of segments ***b*** and ***c***. The corresponding orientation heatmap has three distinct transitions in orientation.

| assembly_accesion | read_type | sequencer(s) | assembler | misassembly_supported | notes |
|---|---|---|---|---|---|
| GCA_005886035.1 | hybrid | Illumina HiSeq 4000, PacBio Sequel | SPAdes | yes | Re-assembly agrees with corrected genome |
| GCA_008065435.1 | long | PacBio RS II | HINGE | yes | Two traversals corresponding to original and corrected genome |
| GCA_009730315.1 | hybrid | Illumina HiSeq 4000, PacBio RS | SPAdes | yes | Two orderings of contigs corresponding to original and corrected genome |
| GCA_016451985.1 | hybrid | Illumina MiSeq, PacBio Sequel | SPAdes | yes | Two ordering of contigs corresponding to original and corrected genome |
| GCA_002012025.1 | short | Illumina MiSeq | SPAdes | n/a | Fragmented re-assembly. Possible read data omitted. |
| GCA_016452025.1 | long | PacBio Sequel | HINGE | n/a | Fragmented re-assembly. Possible read data omitted. |
| GCA_014623225.1 | long | PacBio RS II | n/a | n/a | Manually discarded due to poor heatmap quality |
| GCA_900327275.1 | long | PacBio RS | n/a | n/a | Manually discarded due to poor heatmap quality |
| GCA_003112145.1 | short | Illumina MiSeq | n/a | n/a | Some read data omitted. Re-assembly not possible |
| GCA_003339775.1 | long | PacBio RS | n/a | n/a | Some read data omitted. Re-assembly not possible |
| GCA_000198515.1 | none | | | | |
| GCA_000487155.2 | none | | | | |
| GCA_001562215.1 | none | | | | |
| GCA_001722005.2 | none | | | | |
| GCA_001723625.1 | none | | | | |

| | |
|---|---|
| GCA_001936395.1 | none |
| GCA_001938665.1 | none |
| GCA_002005165.1 | none |
| GCA_002441975.1 | none |
| GCA_004291075.1 | none |
| GCA_009950475.1 | none |
| GCA_009951245.1 | none |
| GCA_011045215.1 | none |
| GCA_011801475.1 | none |
| GCA_012934815.1 | none |
| GCA_013085185.1 | none |
| GCA_013305705.1 | none |
| GCA_014168635.1 | none |
| GCA_014489455.1 | none |
| GCA_014731795.1 | none |
| GCA_900324235.1 | none |
| GCA_900327275.1 | none |

**Suppplementary Table T1.** List of misassemblies detected from 5,000 randomly chosen GenBank genomes.
